# Effect of RNA preservation methods on RNA quantity and quality of field collected avian whole blood

**DOI:** 10.1101/2021.11.16.468897

**Authors:** Johanna A. Harvey, Sarah A. Knutie

## Abstract

A limitation of comparative transcriptomic studies of wild avian populations continues to be sample acquisition and preservation to achieve resulting high-quality RNA (i.e., ribonucleic acids that transfers, translates, and regulates the genetic code from DNA into proteins). Field sampling of wild bird samples provides challenges as RNA degradation progresses quickly and because cryopreservation is often not feasible at remote locations. We collected blood samples from songbirds, as avian blood is nucleated and provides sufficient transcriptionally active material in a small and non-lethal sample, to compare the efficacy of widely available RNA stabilizing buffers, RNAlater (Ambion) and DNA/RNA Shield (Zymo) at differing concentrations along with a dry ice-based flash freezing method (Isopropanol 99% and dry ice mixture, −109°C). Each blood sample was divided among five different preservation treatments (dry ice-based flash freezing, RNAlater with 1:5 or 1:10 dilution, or DNA/RNA Shield with 1:2 or 1:3 dilution). A new protocol was optimized for total RNA extraction from avian blood samples with small starting volumes enabling sampling of small passerines. We quantified quality measures, RNA integrity numbers (RIN^e^), rRNA ratios, and total RNA concentrations. We found that RNA preservation buffers, RNAlater and DNA/RNA Shield at all concentrations, provide sample protection from RNA degradation. We suggest caution against using dry ice-based flash-freezing alone for samples preservation as these samples resulted in lower quality measures then samples in preservation buffer. Total RNA concentration was generally not affected by preservation treatment and may vary due to differences in initial samples volumes and carryover across processing steps.

**LAY SUMMARY:**

- Preserving RNA samples collected in field conditions, under extreme or variable conditions, remains a challenge and continues to limit the study of gene expression in wild birds.
- We optimized RNA extractions for small-volume whole avian blood samples.
- We compared commonly used RNA preservations methods, commercial RNA buffers at varying concentrations along with a dry ice-based flash freezing method, for field collected blood bird samples.
- All RNA buffers preserved samples resulted in high quality RNA extractions suitable for downstream RNA sequencing, while dry ice-based flash freezing resulted in less reliable sample preservation.
- Our results support the feasibility of blood sampling as a method of non-lethal sampling that will enable the increased usage RNA sequencing in wild bird studies.

**KEY TERMS:** *RNA-Seq:* high-throughput sequencing which profiles RNA molecules in a sample at the collection timepoint. Provides content and abundance information and may identify novel genes and isoforms. Prior to sequencing, library preparation may either target mRNA or total RNA with the use of enrichment or depletion steps.

*Transcriptome:* all RNA transcripts, both coding and non-coding, in an individual.

*rRNA ratio (28S/18S):* commonly used ratio of 28S and 18S ribosomal RNA (rRNA) (band sizes or peak area ratio) that informs the integrity of RNA. Appropriate ratios vary by taxonomic groups and associated rRNA sizes. Acceptable ratios of 28S to 18S are approximately 2.0 for eukaryotes.

*Electropherogram:* a quantitative distribution of fragment size.

*Qubit 4 Fluorometer:* fluorescence-based quantification of RNA molecule concentration using fluorescent intercalating dyes which bind to target molecules.

*Agilent 4200 TapeStation:* high-throughput analyzer which uses microfluidics, microcapillary electrophoresis, and fluorescence detection that allows for size determination of the isolated molecules. Used for quality control and quantification of DNA and RNA samples for downstream methods. Reports metrics to assess sample (DNA, RNA, protein) integrity and quantification, which includes: electropherogram, gel-like image, concentration values, 28S/18S rRNA ratios, and an algorithmically determined integrity score (RIN^e^). Provides a more complete characterization of the measured samples then previous analyzers for downstream analysis.

*Fast zone:* area between small RNAs and the 18S rRNA fragment (~200 bp to 1.8 kb). The fast zone is where the degradation of the 18S and 28S peaks accumulates.

*RIN^e^:* RNA integrity number (RIN equivalent) which is determined by quantitative measurements of total RNA degradation. Measurements are based on features of the electropherogram which assesses degradation or lack of degradation by measuring the ratio of area in the fast zone to the 18S peak signal. RIN of 10 signifies the highest quality intact RNA and zero is the worst score indicates a low-quality degraded sample.

## INTRODUCTION

Preservation of ribonucleic acids (RNA) is an essential step in all transcriptomic studies that use field collected samples. RNA-Seq methods allows investigation of genomewide molecular responses to environmental stressors. Transcriptome studies enable the study of quantitative expression, gene fusions, splice variants, novel transcript discovery and examine coding (mRNA) and non-coding sequences (including miRNA, sRNA, and other RNA classes). The accuracy of RNA sequencing results are dependent on the initial preservation of intact RNAs (Wang et al. 2009). Determining proper methods for RNA sample preservation during challenging and variable field collecting conditions is essential to facilitate increased usage of wild RNA-Seq studies, which will improve our understanding of responses to environmental stressors in free-living populations. Field collection of samples is difficult due to the rapid rate of degradation in RNAs and because the options for cryopreservation and accurate temperature control are often not available in remote locations (Cheviron et al. 2011, Chiari and Galtier 2011, Gayral et al. 2011, Camacho-Sanchez et al. 2013). Proper and timely preservation of samples is necessary for accurate and unaltered characterization of the *in vivo* state of cells and associated RNA expression (Gallego Romero et al. 2014).

Ribonucleases (RNases) cause cleavage and rapid degradation of sample RNA which can distort RNA-Seq read coverage, alter gene expression profiles, and cause the loss of rare transcripts (Wang et al. 2016). RNA is fragile due to both the abundance of RNases and its structure which makes it sensitive to hydrolysis, these degradation processes increase the number of small RNAs. Stabilization of RNA samples, halting RNase activity and hydrolysis, requires timely preservation and planning. Immediate freezing in a −80°C or snap-freezing in liquid nitrogen has been considered the gold standard for RNA stabilization and preservation, particularly in clinical and lab settings (Wolf 2013). However, while snap-freezing is still used in controlled experimental settings the use of commercial agents for stabilization has increased greatly. Commercial RNA fixatives, which permeate cells and denature RNases, have previously been found to prevent degradation of RNA and stabilize expression levels (Gorokhova 2005, Cheviron et al. 2011, Vider et al. 2020). These commercial products use proprietary reagents to stabilize RNA: RNAlater, according to its Material Safety Data Sheet, uses ammonium sulfate to stabilize proteins, whereas DNA/RNA Shield, according to a safety declaration, uses guanidine salt which induces protein unfolding. Few studies have tested whether these reagents are effective at preserving RNA from nucleated blood samples collected in the field.

For wild avian studies, blood sampling allows for a non-lethal and minimally invasive method of sample collection facilitating the study of wild avian transcriptomes (Huang et al. 2016). Birds and all non-mammalian vertebrates have nucleated red blood cells, meaning red blood cells as well as leukocytes are transcriptionally active (Chiari and Galtier 2011, Waits et al. 2020). The high quantity of nucleated blood cells reduces the amount of blood needed for transcriptome studies as compared to mammalian studies (i.e., non-nucleated blood cells). The small amount of nucleated blood needed for RNA-Seq can enable studies of critically endangered avian species, species at risk, and small-bodied species.

Blood transcriptomes remain underutilized in free-living avian studies. Transcription is tissue specific making the choice of RNA target material dependent on the research question. However, peripheral blood circulates the body and may inform systemwide gene expression. When specificity of different tissue transcriptomes has been compared, the blood transcriptome demonstrated lower specificity (Bentz et al. 2019), indicating the utility of blood transcriptomes as environmental mediators. We only found nine studies that have used blood as the source material in wild avian studies using RNA-Seq, excluding captive/experimental manipulation studies of wild caught taxa. Blood transcriptomes have been demonstrated to be informative for studies including immunologic, detoxification, lipid metabolism responses as well as recovering a high proportion of genes expressed across the genome and tissue types (Liew et al. 2006, Désert et al. 2016, Höglund et al. 2017, Watson et al. 2017, Bentz et al. 2019). High quality and quantity RNA are necessary for accurate *in vivo* characterization of blood transcriptomes.

Ribosomal RNA (rRNA) makes up approximately 80% of total RNA in eukaryotes, with the remaining total RNA comprised of other RNAs including precursor messenger RNA, messenger RNA (mRNA), and several non-coding RNAs (Muyal et al. 2009). RNA quality and total yield are used to evaluate RNA integrity prior to downstream processing and sequencing. RNA quality is measured by metrics such as RNA integrity number (RIN or RINe) and rRNA ratio (28S/18S). Accepted RIN values vary, most commonly values greater than seven or eight are considered the threshold for RNA-Seq applications (Shen et al. 2018). RIN and the more recently developed RIN^e^ are the current accepted standards for measuring RNA quality and integrity (Schroeder et al. 2006, Botling et al. 2009, Shen et al. 2018). Prior to the development of the RNA integrity numbers rRNA ratios were predominantly used to determine RNA quality. The rRNA ratio for chickens is 2.4 (4.4 kb 28S /1.8 kb 18S; Dyomin et al. 2016). While, complete annotation of the rRNA of passeriform genomes remains unresolved, due to excessive repetition of the rRNA gene cluster, acceptable ratios of 28S to 18S are approximately 2.0 (Dyomin et al. 2016).

To determine best practices for stabilization of whole avian blood from field collected samples, we compared various widely available RNA preservation methods. Specifically, we compared a dry ice-based flash freezing method (Isopropanol 99% and dry ice mixture, −109°C) to two commonly used and commercially available RNA reagent buffers: RNAlater (Ambion, Invitrogen, Carlsbad, CA, USA) at 1:2 and 1:3 concentrations and DNA/RNA Shield (Zymo Research, Irvine, CA, USA) at 1:5 and 1:10 concentrations. To compare preservation methods and dilutions, we isolated total RNA using an optimized extraction protocol and compared measures of quality (e.g., RIN^e^, rRNA ratio) and quantity (e.g., total RNA concentration). The quality and quantity were compared across preservation methods to determine the most successful method of RNA preservation for field collected avian blood samples to be used in downstream RNA sequencing studies.

## METHODS

### Blood sampling and preservation

We captured birds opportunistically with a mist-net in Clearwater County, Minnesota, USA. Individual blood samples (*n* = 8) were collected from three species, including Brown-headed Cowbirds (*Molothrus ater*) (*n* = 3), Red-winged Blackbird (*Agelaius phoeniceus*) (*n* = 1), and Rose-breasted Grosbeaks (*Pheucticus ludovicianus*) (*n* = 4). We collected a total of 100 μL of whole blood from the brachial vein from each individual bird. Specifically, 20 μL of blood was collected into five pre-measured capillary tubes for precise measurement. Each capillary tube of blood was then immediately expelled randomly, using a sterile air blower bulb (VSGO, Shanghai, China), into five different pre-aliquoted treatment cryogenic tubes with different preservation reagents x dilutions. The five treatments included: 1) DNA/RNA Shield 1:2 (20 μL blood + 20 μL DNA/RNA Shield), 2) DNA/RNA Shield at 1:3 (20 μL blood + 40 μL DNA/RNA Shield), 3) RNAlater at 1:5 (20 μL blood + 80 μL RNAlater), 4) RNAlater 1:10 (20 μL blood + 180 μL RNA later), and 5) dry ice + isopropanol bath based flash freezing mixture (−109°C) (20 μL blood only). Samples were then returned to the laboratory at Itasca Biological Station, University of Minnesota, where the samples in RNAlater and DNA/RNA Shield were vortexed for 10 seconds, placed in a 4°C refrigerator for 24 hours, then moved to a −80°C freezer. The dry ice-based flash frozen samples were placed immediately into a −80°C freezer. After one week, all samples were shipped (next day air) on dry ice (−78.5°C) to laboratory facilities at University of Connecticut and stored at −80°C until processed.

### RNA isolation, quantification, and quality assessment

Tri-Reagent based phase separation followed by a column clean-up has previously been found effective for the extraction of total RNA from avian whole blood samples (Chiari and Galtier 2011, Mewis et al. 2014). This two-step extraction method is effective in reducing the high DNA proteins and salt associated with avian blood via the phase separation and the column cleanup. Here we optimized the phase separation and column-based extraction protocol for small avian whole blood samples (~20 μL). RNA extractions were completed in an RNase free environment. A complete step by step isolation protocol can be found at https://github.com/JAHarvey/RNA-Blood-preservation-extraction/blob/main/RNA_Isolation_Protocol.mkd. Total RNA was isolated from 20 μL of peripheral whole blood using a modified Tri-Reagent (Ambion, Invitrogen, Carlsbad, CA, USA) and Direct-zol RNA Miniprep Plus Kit (Zymo Research, Irvine, CA, USA) protocol. The samples were incubated at room temperature for 2 minutes and then centrifuged for 1 minute at 8,000 RCF to lightly pellet the blood. Preservation fluids (i.e., RNAlater, DNA/RNA Shield) were pipetted off leaving no more than ~15 μL of preservative and 500 μL of Tri-Reagent were added along with a sterile 5mm stainless steel bead (Thomson, Radford, VA, USA) the samples were then vortexed for 30 seconds before adding an additional 500 μL of Tri-Reagent. The sample was then vortexed for 10 minutes at room temperature. The phase separation portion of the Tri-Reagent protocol was then followed, and the upper aqueous phase (500 μL) was transferred to new microcentrifuge tubes. We then used column purification with the Direct-zol RNA Kit, following manufacturer’s protocol beginning with the RNA purification step and including DNase treatment. We eluted total RNA using 50 μL of RNA/DNA free water. We quantified total RNA concentration (ng/μL) using a Qubit RNA HS assay kit on a Qubit 4 Fluorometer (Thermofisher Scientific, Waltham, MA, USA). We used a 4200 TapeStation Analyzer System and High Sensitivity RNA ScreenTape assays (Agilent, Santa Clara, CA, USA) to determine rRNA ratio, total RNA concentration, and RNA integrity numbers (RIN^e^).

### Statistical Analysis

Generalized linear mixed models (GLMMs) were used to determine the effect of different preservation treatments on total RNA quantity (Tapestation and Qubit concentration) and quality (RIN^e^, rRNA ratio ratio), with individual bird as a random effect. When the response variables did not pass the Shapiro-Wilk normality test, they were log transformed for the analyses. GLMM analyses were carried out using the lmer function with the lme4 package (Bates et al. 2015) and the ANOVA function with the car package (Fox and Weisberg 2011). Pairwise post-hoc comparisons were compared among treatments using Tukey’s HSD (honestly significant difference) tests in the package emmeans (Lenth et al. 2018). Pearson r correlations were used to determine the strength of the relationships among measures of RNA quality and quantity. All statistical analyses were conducted, and figures were created, in R version 4.1.0 and RStudio (v1.4.1717).

## RESULTS

Preservation of samples at the time of collection is critical in all downstream sample preparation and sequencing steps. We found that preservation treatment affected both quality metrics (Table 1), RIN^e^ (Fig. 1A; χ^2^ = 37.59, df = 4, *P* = < 0.0001) and rRNA ratio (Fig. 1B; χ^2^ = 19.50, df = 4, *P* = 0.0006). For RIN^e^, RNAlater had the highest values and the lowest variance (mean ± SD: 9.59 ± 0.14, Table 1), while dry ice-based flash frozen samples had the lowest scores (mean ± SD: 8.36 ± 0.84, Table 1). RIN^e^ for dry ice-based flash frozen samples were significantly lower than samples preserved in preservation buffers at all concentrations (Tukey post-hoc test, *P* < 0.01). Differences among RNA preservation buffer treatments were not significant (Tukey post-hoc test, *P* ≥ 0.57). For rRNA ratio, the samples preserved in RNAlater at a 1:10 concentration had significantly higher values (mean ± SD: 1.71 ± 0.15, Table 1) than the dry ice-based flash frozen samples (mean ± SD: 1.18 ± 0.40, Table 1; Tukey post-hoc test, *P* = 0.004). No other differences among treatments were significant for rRNA ratio samples (Tukey post-hoc test, *P* ≥ 0.17)

**Table 1.**
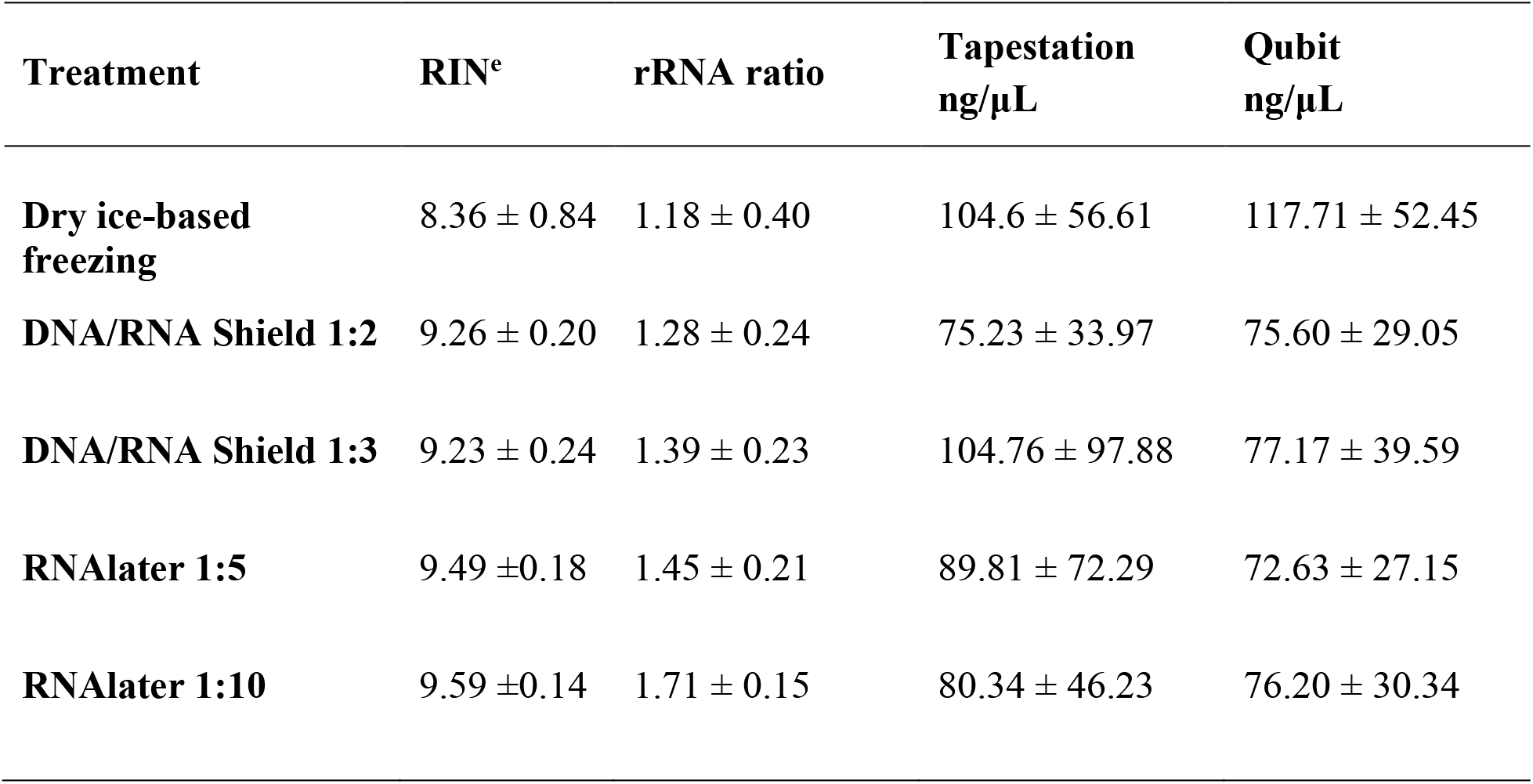
Average (mean ± standard deviation) of quality metrics, RIN^e^ and rRNA ratio, and total RNA concentrations as measured by the Tapestation and Qubit.

| Treatment | RIN<sup>e</sup> | rRNA ratio | Tapestation<br>ng/μL | Qubit<br>ng/μL |
| --- | --- | --- | --- | --- |
| **Dry ice-based freezing** | 8.36 ± 0.84 | 1.18 ± 0.40 | 104.6 ± 56.61 | 117.71 ± 52.45 |
| **DNA/RNA Shield 1:2** | 9.26 ± 0.20 | 1.28 ± 0.24 | 75.23 ± 33.97 | 75.60 ± 29.05 |
| **DNA/RNA Shield 1:3** | 9.23 ± 0.24 | 1.39 ± 0.23 | 104.76 ± 97.88 | 77.17 ± 39.59 |
| **RNAlater 1:5** | 9.49 ± 0.18 | 1.45 ± 0.21 | 89.81 ± 72.29 | 72.63 ± 27.15 |
| **RNAlater 1:10** | 9.59 ± 0.14 | 1.71 ± 0.15 | 80.34 ± 46.23 | 76.20 ± 30.34 |

**Fig. 1.**
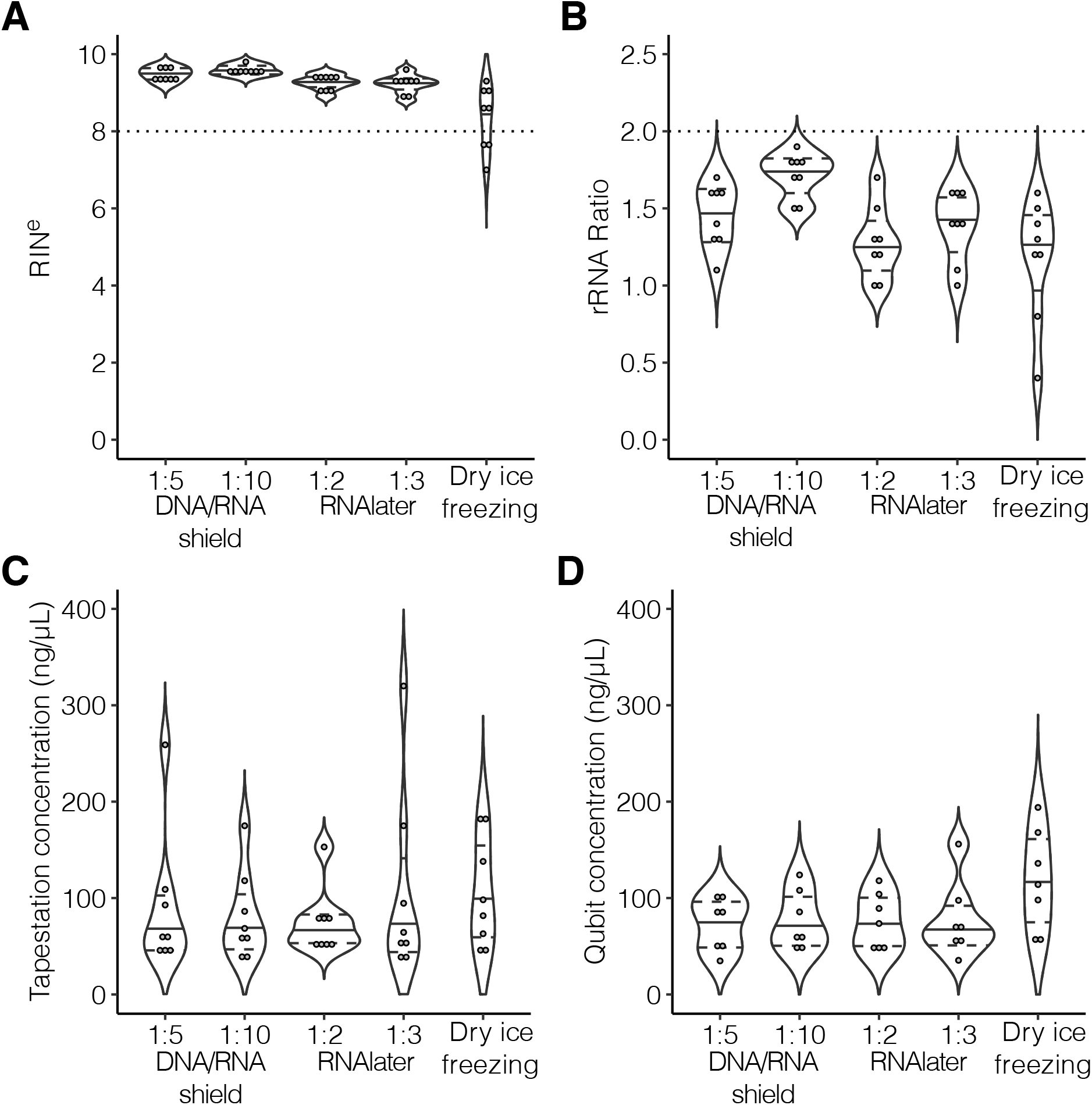
Effect of treatment on A) RIN^e^, B) rRNA ratio, C) Tapestation RNA concentration (ng/μL), and D) Qubit RNA concentration (ng/μL). Each point represents an individual sample.

We found that preservation treatment had little effect on quantity metrics; differences in Qubit concentrations were marginally significant (Fig 1D; χ^2^ = 11.04, df = 4, *P* = 0.03) but there was no significant effect for Tapestation concentrations (Fig. 1C; χ^2^ = 4.40, df = 4, *P* = 0.36). Qubit RNA concentrations from the dry ice-based flash frozen samples were higher (117.71 ± 52.45, Table 1) than samples across all preservation treatments but difference among preservation buffer treatments were not significant (Tukey post-hoc test, *P* ≥ 0.09). Both quantity measures, Tapestation and Qubit, total RNA concentrations were highly correlated (Fig. 2B; Pearson’s r = 0.90) to each other, as were quality measures RIN^e^ and rRNA ratios (Fig. 2A; Pearson’s r = 0.73). However, neither quality value was related to either RNA quantity measure (Pearson’s r < 0.30 for all pairs).

**Fig. 2.**
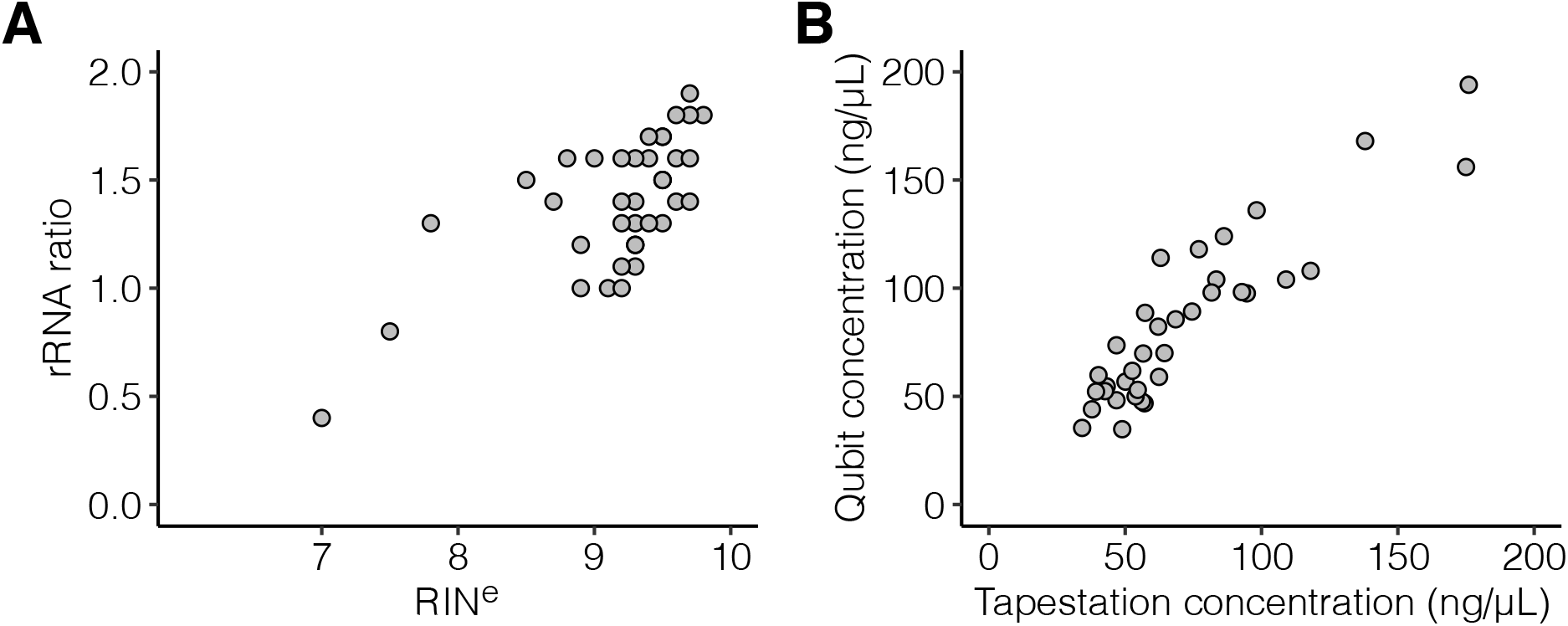
Relationships between the A) RIN^e^ and rRNA ratio (quality), and B) Tapestation and Qubit RNA concentrations (quantity).

## DISCUSSION

Non-lethal sampling of peripheral avian blood provides sufficient transcriptionally active RNA enabling genomewide transcriptome studies of at risk and endangered species. Here we provide a comparison of RNA preservation methods for field sampled whole avian blood. We tested preservation methods in widely available commercial buffers at various concentrations as well as a dry ice-based flash freezing method in place of liquid nitrogen flash freezing. Both preservations buffers, RNAlater and DNA/RNA Shield, at all concentrations, produced extractions with high quality scores demonstrating that RNA preservation buffers, successfully preserved RNA from whole avian blood. RNAlater at 1:10 concentrations did recover the highest RIN^e^ and rRNA ratios, both with the lowest variance across treatments. We caution that handling time of samples prior to preservation as well as during downstream processing may increase degradation and increase bias of *ex vivo* effects on results. Additionally, we did not test extended storage periods of RNA preserved samples at ambient or field temperatures as this has been found to lower RNA quality and impact expression profiles (Camacho-Sanchez et al. 2013, Passow et al. 2019). Our goal was to test RNA preservation methods for storage conditions commonly available to us in the field, conditions where liquid nitrogen is not available but cold storage is available. Often field-sampling conditions do not have cold storage immediately available, but 4°C can be achieved with a cooler and ice packs, and thus reagent preservation is necessary.

While flash freezing with liquid nitrogen (−190°C) is considered the standard for RNA preservation (Wolf 2013), liquid nitrogen was not available at the field site and is commonly difficult to acquire at remote sites when field sampling. We expected our dry ice-based flash freezing method (−109°C) to provide a suitable substitute for liquid nitrogen flash freezing and be able to serve as the standard comparison for comparing the commercial stabilization buffers. Unfortunately, the dry ice-based flash freezing method performed less reliably. The dry ice-based flash frozen samples had lower and more variable quality scores than samples collected in either of the preservative buffers at any concentration. More than half of the RIN^e^ values for the dry ice-based flash freezing method are still suitable for downstream RNA-Seq as they meet the commonly accepted threshold of RIN > 8. Previous comparisons of methods have shown that flash freezing tissue samples in the field with liquid nitrogen yielded comparable quality RNA extractions and higher RNA concentrations as compared to RNAlater (Camacho-Sanchez et al. 2013, Passow et al. 2019). The performance of this dry ice-based flash freezing method may have been impacted by sublimation rate of dry ice at high ambient temperature. Alternatively, the increased degradation may be due to uneven freezing of the blood samples. This method of dry ice-based flash freezing, or the use of dry ice alone are not recommended for field collection and preservations of avian blood RNA samples alone without the use of preservation buffers.

Lower rRNA ratios, re indicate higher degradation has occurred in the sample as the 28S peak is known to decrease faster than the 18S peak. However, only weak correlations of RNA quality and rRNA ratios have been previously found (Imbeaud et al. 2005), indicating the measure of rRNA ratio alone, though informative, may be an unreliable metric of mRNA quality. RNA integrity number (RIN and RIN^e^) is preferentially used to assess RNA quality prior to downstream sequencing, as it provides the most complete characterization of RNA quality by measuring features of degradation (see Schroeder et al. 2006).

Tapestation and Qubit concentrations (ng/ μL) for all samples were sufficient for downstream sequencing. However, total concentration measures varied across analyzers. The Qubit measures florescence via target-specific dyes, while the Tapestation calculates the concentrations using the cumulative area of electropherogram sample peaks (28S and 18S) compared to the lower marker, which is a peak of standardized and calibrated concentration. Quantity measurements assess total RNA concentrations, this measurement includes many small RNAs and when measured with a fluorometer this measurement may not be able to differentiate degraded RNA. Though the amount of RNA is somewhat indicative of quality, this measurement varies by starting input material, carryover across extraction steps, and is expected to vary when measured with different quantification methods such as the Qubit Fluorometer and the Tapestation. Concentration of RNA extracted from avian and reptile blood (both nucleated blood cells) have been found to vary in concentrations despite the equal starting inputs for the extraction (Chiari and Galtier 2011). These differences in concentration may be due to differences in hydration levels of sampled individuals, upon blood draw the blood may appear thinner or thicker. Overall, total RNA quantity was generally not affected by preservation treatment and is not as important of a limitation to downstream usage as RNA quality.

The next step could be to create sequencing libraries and sequence the samples to determine specific differences across methods. Library preparations may further impact the downstream sequencing results of peripheral avian blood and the selection of appropriate methods varies with the intended objective of sequencing. These options include removal of ribosomal RNA (rRNA) which comprises 90% of total RNA (Wilhelm and Landry 2009). Removal of rRNA may be carried out via polyadenylated region selection, which targets transcripts with polyA tails present as in most mRNAs, or via rRNA depletion which binds or targets rRNA for subsequent removal. However, this is beyond the scope of the study.

## Conclusion

Here, we demonstrate that field collection of high quality preserved peripheral avian blood RNA samples is possible with available commercial buffers. Our results will enable future study of avian blood transcriptomes in wild avian populations. To this day, most avian transcriptome studies have been conducted in clinical and laboratory settings. Controlled experimental setups in captive species are informative and allow researchers to reduce the number of variables impacting study species. However, captive studies do not capture the complex transcriptomic response of wild species in dynamic natural environments as the transcriptomic response has accurately been described as a “snapshot” of cellular response to its environment (Alvarez et al. 2015). While we only examined preservation of avian blood samples for RNA extraction, these preservation methods are applicable to other taxa with nucleated blood cells, i.e., amphibians, reptiles, and fish. The increased implementation of RNA-Seq studies in wild birds via non-lethal blood sampling will help inform wild avian transcriptomic responses to environmental stressors.

## ACKNOWLEDGEMENTS

We thank Itasca Biological Station, including Lesley Knoll and Laura Domine, for research and logistical support. We thank Helen and Ken Perry for allowing us to catch birds on their property and Alexandra Parker and Lauren Albert for assistance in collection of field samples. We would like to thank Jonathan Puritz, Spencer Galen, and Kristi Sikkink for advice on extraction methods.

## Author contributions

J.A.H. and S.A.K. conceived the idea, design, experiment; S.A.K. collected field data and edited the paper, J.A.H. conducted the research and wrote the paper, J.A.H. and S.A.K. analyzed the data. Both authors approved the manuscript before final submission.

